# Vision does not impact walking performance in Argentine ants

**DOI:** 10.1101/2020.05.05.079582

**Authors:** G.T. Clifton, D. Holway, N. Gravish

## Abstract

Many walking insects use vision for long-distance navigation, but the influence of vision in detecting close-range obstacles and directing the limbs to maintain stability remains largely untested. We compared Argentine ant workers in light versus darkness while traversing flat and uneven terrain. In darkness, ants reduced flat-ground walking speeds by only 5%. Similarly, neither the approach speed nor the time to cross a step obstacle was affected by lighting. To determine if tactile sensing might compensate for vision loss, we tracked antennal motion and observed shifts in spatiotemporal activity due to terrain structure but not illumination. Together, these findings suggest that vision does not impact walking performance in Argentine ant workers. Our results help contextualize eye variation across ants, including subterranean, nocturnal, and eyeless species that walk in complete darkness. More broadly, our findings highlight the importance of integrating vision, proprioception, and tactile sensing for robust locomotion in unstructured environments.

## Introduction

Walking involves long-distance navigation, avoidance of intermediate-range obstacles, and coordination of the body to ensure stability on uneven terrain (Logan et al., 2010) (Fig. 1). In many insects, such as ants, navigation relies on visually sensing distant environmental features (Cheng et al., 2006; Graham and Cheng, 2009; Graham and Philippides, 2017; Narendra et al., 2013b; Wehner and Muller, 2006). At a close range, vision informs slow reaching of the limb, such as during gap crossing (Niven et al., 2010; Pick and Strauss, 2005). However, insect walking involves rapid limb movements that may exceed visual sensory delays, possibly precluding vision-based responses to obstacles and terrain features (Haselsteiner et al., 2014; Höltje and Hustert, 2003). To test how vision impacts walking performance, we tracked ant walking speeds and step obstacle crossing in light and dark conditions.

**Figure 1.**
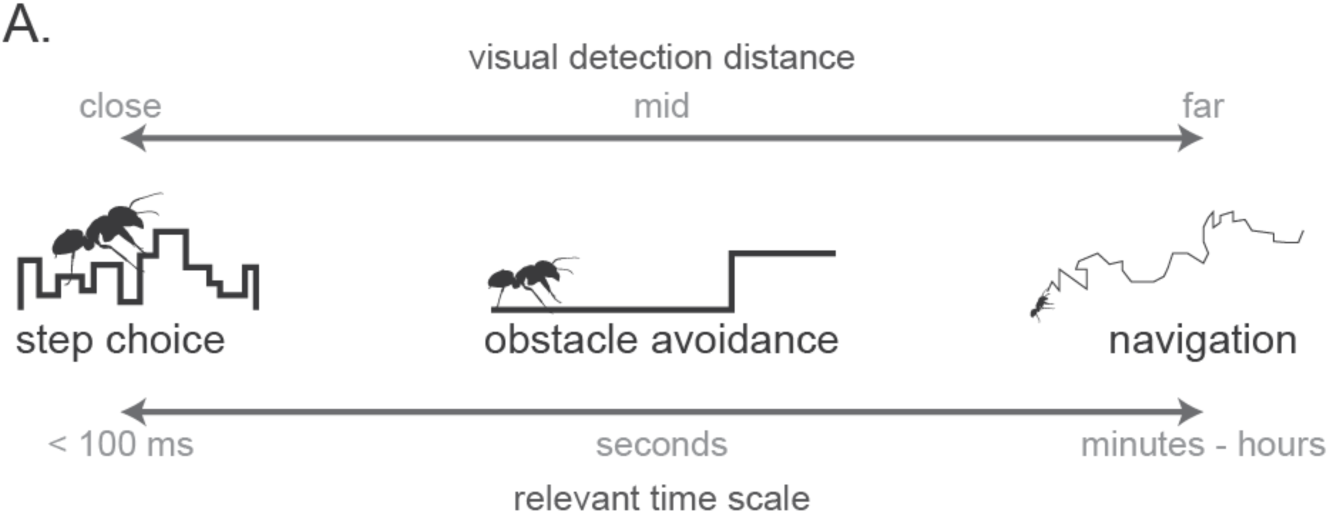
Impact of vision on ant walking across spatiotemporal scales. (A) Ants navigate by sensing long-distance visual features, however visual perception of mid- or close-range objects may influence walking performance. Closer-range visual perception restricts effective feedback timing, with fast step frequencies potentially preventing ants from using visual perception at these scales.

Ants walk by taking quick steps, which may preclude visual control observed during slower limb motions. Like other hexapedal insects, ants typically walk by stepping with three limbs at a time in an alternating tripod gait (Zollikofer, 1994). This gait is most often associated with fast walking insects like cockroaches (Full and Tu, 1990) as opposed to slow, limb-by-limb “metachronal” movements in ambling walkers such as stick insects (Graham, 1972). Although recent studies identify considerable variation between these extreme gaits (Bender et al., 2011; Szczecinski et al., 2018), comparing slow, stick-insect-like walking versus fast, cockroach-like walking highlights constraints in the neural control of these motions. Slow movements of the limb often incorporate information from visual signals (Blaesing, 2004; Collett, 2002; Dürr, 2001; Niven et al., 2010; Niven et al., 2012; Pick and Strauss, 2005). This closed-loop control substantially guides walking in large animals (Wilkinson and Sherk, 2005) and humans (Patla and Greig, 2006; Reynolds and Day, 2005a; Reynolds and Day, 2005b; Smid and den Otter, 2013). However, as body size decreases there are increases in both step frequency (Heglund et al., 1974; Lee et al., 2016) and substrate unevenness (Kaspari and Weiser, 1999), furthering the need for secure foot placement but limiting the reflex time for visual sensory signals.

The ability for vision to inform locomotion planning requires both visual detection of environmental features and rapid processing of this information. Transduction speeds for photoreceptors range from 30-150 ms in cockroaches (Heimonen et al., 2012; Ignatova et al., 2020), 35-60 ms in hawkmoths (Krishnan and Sane, 2014) and 30-250 ms in blowflies (Land and Collett, 1974; Warzecha and Egelhaaf, 2000). Comparatively, neural conduction speeds are relatively fast at 0.5-3.7 m/s in cockroaches (Pearson et al., 1970). Ants walk at an average of 10-12 strides per second (Clifton et al., 2020; Reinhardt and Blickhan, 2014), limiting intra-stride response times to within 100 ms. Given the relatively slow transduction speeds of walking insect photoreceptors (Frolov et al., 2017), fast movements of the limbs during walking may preclude visual feedback (Full and Koditschek, 1999). Nocturnal and subterranean species particularly confront feedback constraints since transduction speeds depend on both light levels and temperature (Frolov and Ignatova, 2020; Heimonen et al., 2012; Warrant, 2017; Warzecha and Egelhaaf, 2000). These data suggest that vision likely does not influence intra-stride walking coordination but may inform obstacle avoidance at intermediate distances. However, few studies have directly tested the impact of vision on step-to-step walking performance with “effective blindness” presented as a novelty (Gilbert, 1997; Jayaram et al., 2018), whereas it may be considerably widespread.

In addition to sensing their surroundings using vision, many walking insects rely on chemosensory and tactile information. Insect antennae span diverse morphologies, sensing chemicals, heat, humidity, and mechanical feedback (Krishnan and Sane, 2015). During walking both cockroaches and stick insects sweep their antennae to identify nearby obstacles (Baba et al., 2010; Dürr et al., 2001; Harley et al., 2009; Okada and Toh, 2004; Okada and Toh, 2006). Ants possess jointed antennae that they actuate to identify odors (Draft et al., 2018) and tactile features (Klotz and Reid, 1992) along foraging trails. Ant antennae also mechanically jam along tunnel walls to prevent workers from falling (Gravish et al., 2013) and to help in conspecific discrimination (Ozaki et al., 2005; van Zweden and d’Ettorre). However, to the best of our knowledge, antennal activity has not been directly associated with ant walking performance or compared across lighting conditions. Mechanosensory feedback is faster than visual feedback (Sherman and Dickinson, 2004; Yarger and Fox, 2016) However, mechanical feedback stems from both antennae and the limbs (Bingman et al., 2017; Hebets, 2002; Isakov et al., 2016), which may differentially contribute under varying environmental conditions.

The Argentine ant (*Linepithema humile*) forages both diurnally and nocturnally as conditions permit (Abril et al., 2007; Human and Gordon, 1996) with workers regularly walking up to 60 m away from their nests (Hogg et al., 2018; Vega and Rust, 2003). Pheromone-based recruitment trails often cross a variety of substrates, including uneven terrain. Workers possess relatively large eyes compared to closely-related species (Wild, 2004), but, unlike most other species (Knaden and Graham, 2016), visual cues do not impact navigation (Aron et al., 1993). The Argentine ant’s ability to travel relatively large distances in both light and dark conditions makes this species an appealing model for studying how vision impacts walking performance.

Effective walking is integral to the survival of Argentine ant colonies. Thus, one would expect selective pressures to optimize sensory feedback towards maintaining foraging performance under varying environmental conditions. However, vision does not contribute to long-distance navigation in this species (Aron et al., 1993) and the latency of visual reaction times in invertebrates may prevent intrastride feedback, which is particularly relevant for identifying stable footholds on uneven terrain. Instead, we expect vision to impact the identification of and preplanning for close-range obstacles. We test the influence of vision by tracking full-body walking speeds on flat versus uneven terrain and in bright versus dark conditions. Since antennal activity could compensate for lost visual feedback in darkness, we also quantify and compare antennal activity. We show that darkness causes a relatively minor decrease in walking speed compared to terrain unevenness, with no measurable difference in step obstacle crossing or antennal activity. Vision overall plays a minor role in Argentine ant walking.

## Results and Discussion

### Argentine ant eye anatomy

The compound apposition eyes of insects are comprised of ommatidia, which each include a lens and multiple receptors. Quantifying the visual acuity of compound eyes requires measurements of both external features (e.g. ommatidia number, facet dimensions) and internal features (e.g. rhabdom dimensions) (Land, 1997a). Argentine ant workers possess between 80-100 ommatidia per eye (Wild, 2004) with an eye area of 0.026 mm^2^ and an interommatidial angle of 15.3 degrees. Ommatidia facets are approximately hexagonally-spaced with areas ranging from 184-302 µm^2^ and diameters ranging up to 20-26 µm (Figs. 2, S1). Medial and lateral facets were relatively smaller, with the largest facets along the anterior and posterior margins of the eye (Fig. 2).

**Figure 2.**
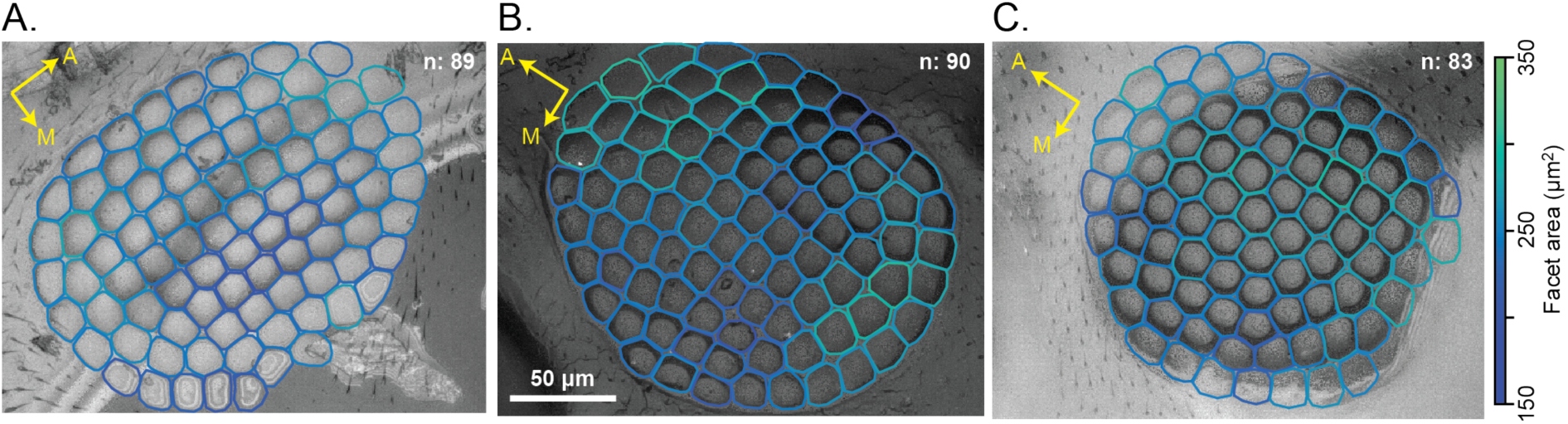
Facet outlines of the eyes of three Argentine ant workers. Facets were manually outlined with colors to represent facet area. The total number of facets for each eye is listed in the panel, however not all facets were able to be outlined. The arrows show the anterior and medial orientations of the head. The scale bar shown in panel B applies across the figure.

The eyes of Argentine ant workers have fewer ommatidia compared to other species that have been studied: *Melophorus bagoti* with 590 (Schwarz et al., 2011), *Polyrhachis sokolova* with 596 (Narendra et al., 2013a), *Camponotus pennsylvanicus* with 375-660 (Klotz et al., 1992), *Formica integroides* with 700 (Bernstein and Finn, 1971), *Camponotus consobrinus* with 800 (Narendra et al., 2016), *Cataglyphis bicolor* with 1200 (Menzi, 1987), four *Myrmecia* species with 2300-3600 (Greiner et al., 2007; Klotz et al., 1992), and *Gigantiops destructor* with 4100 (Gronenberg and Hölldobler, 1999). However, vision-based navigation does not require high resolution (Milford, 2013; Stürzl et al., 2015) and ants with as few as 72-80 ommatidia alter foraging patterns in response to landscape variation (Pratt et al., 2001). Therefore, Argentine ants with 80-100 ommatidia likely can sense long-range visual cues despite their observed disregard of these cues while foraging (Aron et al., 1993). Three species (*Solenopsis invicta, Solenopsis richteri*, and *Temnothorax rugatulus* (Baker and Ma, 2006; Ramirez-Esquivel et al., 2017)) possess a similar number of ommatidia compared to the Argentine ant, however in all cases individual facets are smaller than those of Argentine ant workers. Optical sensitivity is a complex trait that depends on numerous factors, but larger facets increase light capture and are often associated with nocturnal foraging (Greiner, 2006; Sheehan et al., 2019). With relatively large facets at the anterior and posterior margins of the eye, the Argentine ant may benefit from improved vision along the ground and skyline.

### Darkness minimally influences walking speeds on flat and uneven terrain

Five experimental “colonies” of Argentine ants established pheromone-based recruitment trails through a tunnel with a series of observation regions presented in randomized order: 1) flat ground, 2) an uneven checkerboard, 3) a low-contrast single step, and 4) a high-contrast single step (Fig. 3A-B). High-speed videos were recorded across illumination conditions (4204 videos, 240Hz framerate, 3 seconds each). Ants were observed in either visible light conditions or dark conditions imaged with infrared background and flood lighting (Fig. 3C). While the spectral sensitivity of Argentine ant compound eyes has not been directly tested, the photoreceptors of all tested ant species (and Hymenopterans more broadly) are insensitive to light with wavelengths exceeding 650 nm (Aksoy and Camlitepe, 2018; Briscoe and Chittka, 2001). Thus, our “dark” condition illuminated by 850 nm infrared light (<1.8 lux) is likely not perceptible to Argentine ants. While the light conditions presented here (200-2,500 lux) reflect daylight conditions, nine additional colonies were analyzed under dusk light conditions (∼60 lux). The dusk light findings are consistent with those presented here and can be found in the Supplementary Information (Figs. S10-14).

**Figure 3.**
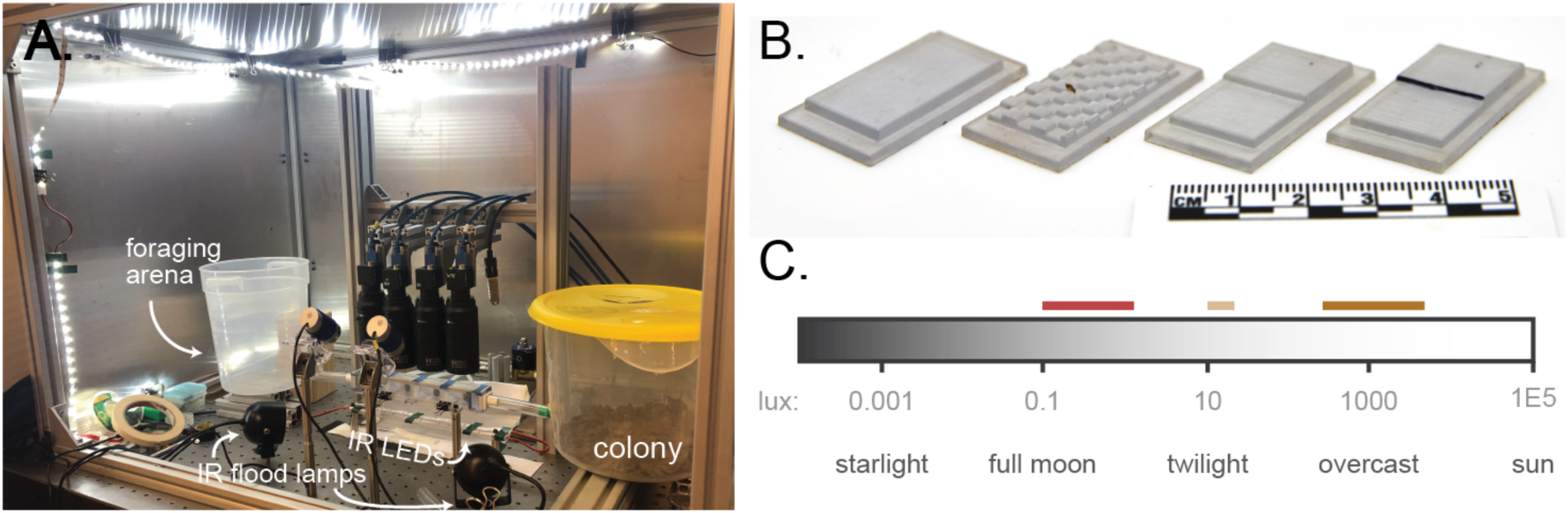
Experimental set-up for quantifying walking performance with varying lighting. (A) A fully-enclosed arena housed a colony of Argentine ants. Workers formed a pheromone trail through a 3D-printed tunnel connected to a foraging arena. Four high-speed machine vision cameras recorded ant motion in the tunnel. White LEDs, infrared flood lights, and infrared strip LEDS beneath the tunnel illuminated the enclosure. (B) Four 3D-printed substrates were inserted in random order into the tunnel floor, including a flat control, a 3 mm checkerboard uneven substrate, and two step obstacles. (C) Light levels vary by over 10^8^-fold throughout the day. Argentine ants walk in complete darkness while in tunnels and forage throughout the day and night. We measured ant walking performance under dark (red), bright (yellow), and dusk (light yellow, results in Supplemental Information) conditions. Lux values obtained from (Cronin et al., 2014)

On flat ground under illumination, ants preferred to walk at a forward speed of 19.2 mm/s, with 5% of the distance traveled occurring at speeds above 28.4 mm/s (Fig. 4C, S3). In darkness, preferred speed reduced by 4.7%, with a median speed of 18.3 mm/s (Mann-Whitney U test, p < 0.001). This pattern was consistent across all colonies except one, which exhibited increased speed in the darkness (Fig S4). Ants slowed down considerably on the checkerboard substrate under light conditions, with a median speed of 11.6 mm/s, 39.6% of the preferred speed under flat/light conditions. Compared to this large decrease, preferred speed on uneven substrates declined only by 1.1 mm/s in darkness (9.5%; Mann-Whitney U test, p < 0.001). Because these speed distributions represent the forward speed aligned with the ant’s orientation, shifts in speed preference could derive from changes in turning or trailway shape. We compared the tortuosity of paths under all conditions using measures of straightness, sinuosity, and fractal dimension (Fig. 4D, S4) (Almeida et al., 2010; Benhamou, 2004; Nams, 1996). While paths on the uneven substrate were less straight than on flat ground (Table S2), there was no influence of lighting (p = 0.33-0.85) (Table S3).

**Figure 4.**
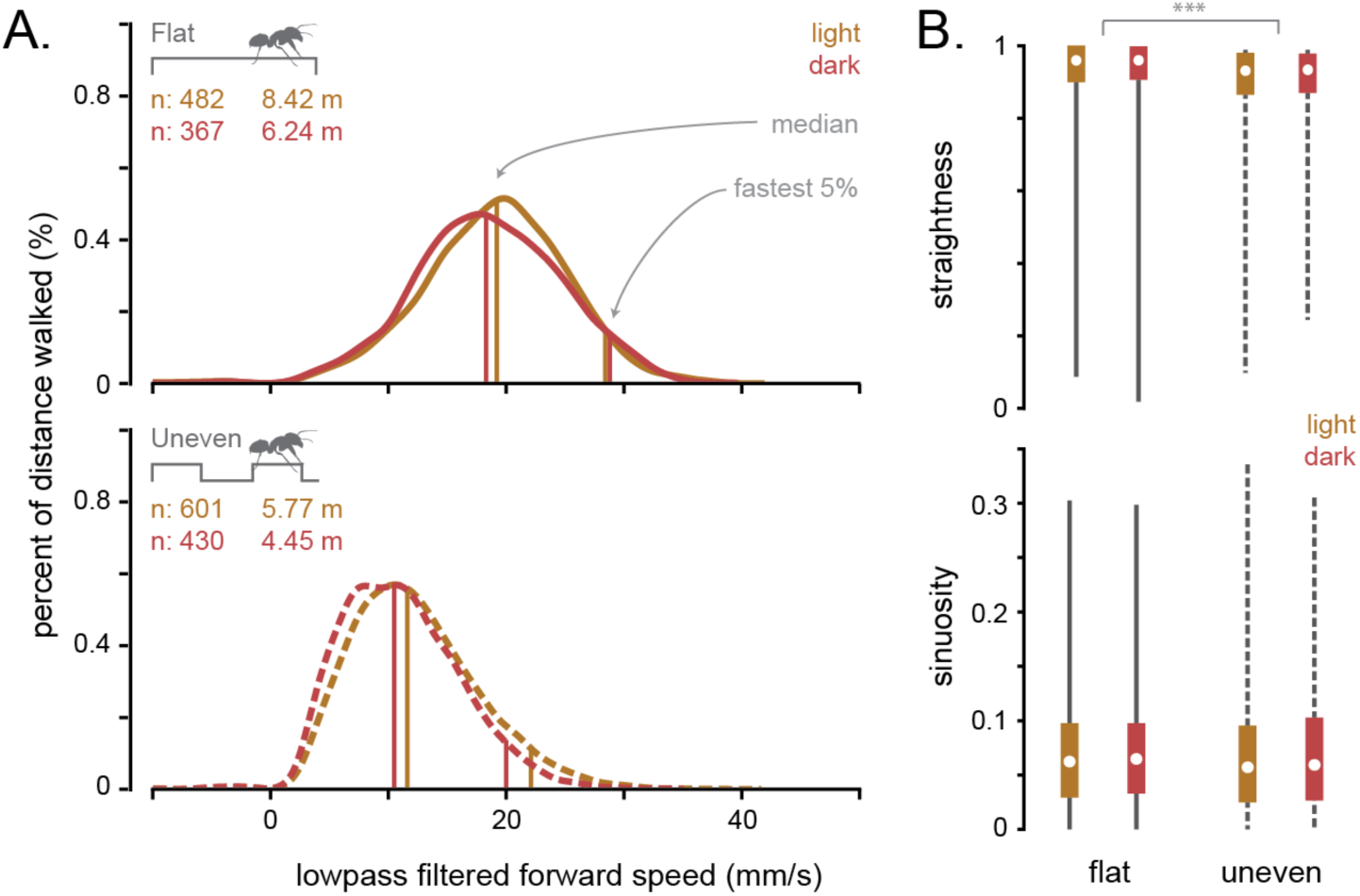
Preferred walking speed and tortuosity on flat and uneven substrates. (A) Distributions showing the distance traveled across speeds for ants on flat (above) and uneven (below) substrates in light (yellow) and dark (red). Vertical lines represent the median speeds and the cutoff for the fastest 5% speed. Light versus dark distributions were significantly different on both flat and uneven substrates (Mann-Whitney, p < 0.001) but with a relatively small effect size (4.7% and 9.5% respectively). (B) Measures of straightness and sinuosity of trackways on flat (solid lines) and uneven (dashed lines) terrain in light (yellow) and dark (red) conditions. Boxes represent interquartile range with whiskers spanning all data points without excluding outliers. Substrate type significantly, but minorly, impacted straightness (Mann-Whitney, p < 0.001), but lighting had no influence on either metric (see Table S2-S3).

We find that Argentine ant walking speeds were minimally impacted by darkness with a < 5% overall reduction on flat ground and with a speed increase for one colony. This finding differs from walking observations in both cockroaches (Baba et al., 2010; Ye et al., 2003) and blowflies (Kress and Egelhaaf, 2012), which slow down by 35% and > 50%, respectively, in darkness. Although fruit flies use optic flow to regulate walking speed (Creamer et al., 2018), they mostly maintain normal speeds in darkness (5-8% reduction) (Howard et al., 2019). To the best of our knowledge, walking speeds in ants have been associated with light levels only in the bull ant, *Myrmecia piriformis*, which showed a logistic increase in speed with luminance (Narendra et al., 2013b). Ant walking speed likely contributes to colony-level survival because it increases rates of food acquisition (Fewell, 1988) and decreases risks associated with foraging, such as predation (Hurlbert et al., 2008; Jayatilaka et al., 2011) and desiccation (Schilman et al., 2005). Therefore, nocturnal foraging in Argentine ants likely does not incur these costs. Interestingly, we observed a greater influence of darkness on speed for walking on uneven versus flat terrain. This finding suggests that ants may rely more on vision when negotiating challenging or uncertain terrain, highlighting the importance of studying locomotion and behavioral responses on non-flat terrain.

### Darkness does not impact crossing a step obstacle

The checkerboard substrate tested above introduces continuous, short-range unevenness. While darkness does not appear to influence the ability of Argentine ant workers to cross this continuously varying terrain, vision may still help identify obstacles at a mid-range distance and to induce pre-planning. Comparisons to other ant species suggest that Argentine ant workers, which possess only 80-100 ommatidia per compound eye, likely cannot identify mid-range obstacles (Palavalli-Nettimi and Narendra, 2018). To test whether workers respond to a looming obstacle, we compared their performance crossing a single 1-mm step in light and dark conditions. Step obstacles are common in perturbation studies (Birn-Jeffery et al., 2014; Gart and Li, 2018; Harley et al., 2009; Theunissen et al., 2015; Watson et al., 2002), and are often used to test neuromechanical control under new conditions. A step height of 1 mm approximates worker coxa height therefore inducing a step and not vertical climbing, which shifts walking kinematics (Weihmann and Blickhan, 2009). Given an interommatidial angle of ∼15 degrees, Argentine ants should be able to visually resolve the step at a distance of at least 3.49 mm (Land, 1997b). We tested step crossing with a low-contrast step, and a high-contrast step, both in light and dark conditions to determine if worker ants adjust their walking kinematics prior to reaching the step.

Argentine ant workers approached the step at an average speed similar to that found on flat substrates (Fig. 5A, solid horizontal line). Workers decelerated when the mandible reached approximately 2 mm from the step (Fig. 5A). While crossing the step, forward velocity remained low and then returned to the average flat ground speed once the worker had moved beyond 5 mm beyond the step. The decrease in speed when crossing the single step was consistent with the preferred speed on the uneven checkerboard substrate of the same step height (Fig. 5A, dashed horizontal lines). This finding suggests that the checkerboard substrate we used may act as a series of step obstacles, each requiring an associated deceleration (Fig. S6).

**Figure 5.**
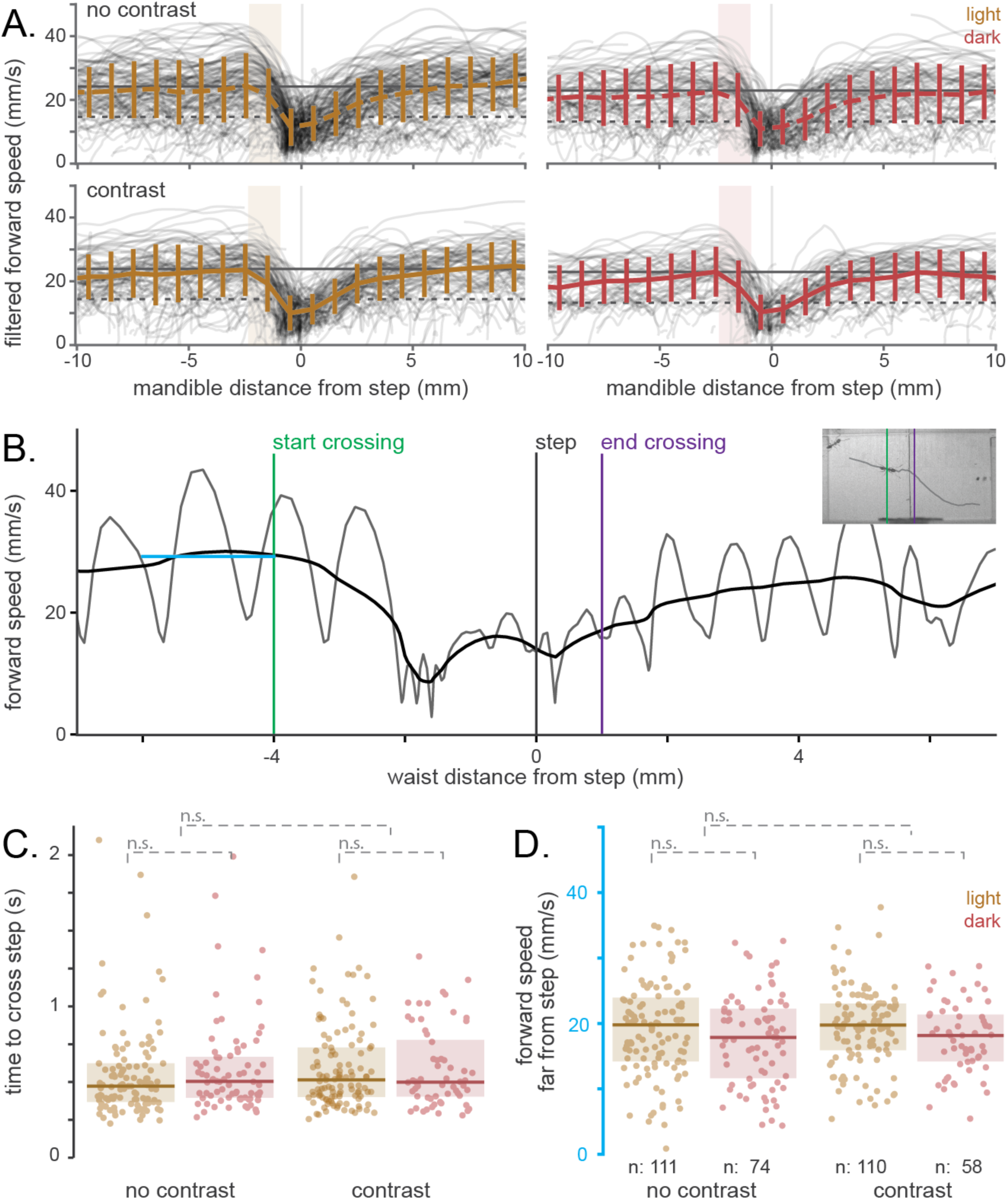
Performance of Argentine ant workers crossing a 1-mm step obstacle. (A) Low-pass filtered forward speeds did not decelerate until the ant’s antennae reached the step edge in both light (yellow) and dark (red) conditions. Vertical bars represent +/-1 standard deviation of all speeds observed in 1-mm windows. Horizontal lines represent preferred, median speeds observed on flat (solid) and uneven (dashed) substrates. Transparent red and yellow boxes represent the normal location of the antennal tips in front of the mandible (1-2.3 mm). (B) Step crossing was defined as starting when the ant waist reached 4 mm from the step (green line) and continued until crossing 1 mm after the step (purple line). Low-pass filtering the speed removed the relatively large accelerations within each step. The median forward speed far from the step (blue dashed line) was determined from when the ant waist was between 6 and 4 mm in front of the step. (C) The time to cross the step did not depend on substrate type (chi-squared LRT test, p = 0.30) or lighting (chi-squared LRT test, p = 0.82). Boxes show the median and interquartile range of the distributions. (D) The forward speed far from the step does not differ for substrate (p = 0.73) or lighting (p = 0.07) conditions.

To reliably compare step crossing performance in all conditions, we defined step crossing as starting at a distance of 4 mm from the step (before any observed deceleration) and ending when the petiole of the worker passed 1 mm beyond the step (Fig. 5B). The time required for workers to cross the step did not differ between light and dark conditions (LME chi-squared test, p = 0.82) or between high-contrast and low-contrast steps (LME chi-squared test, p = 0.30) (Fig. 5C). Despite a 6.5% difference in walking speed far from the step (Fig 5D; LME chi-squared test, p = 0.07), speeds close to the step did not differ (Fig. S7C; LME chi-squared test, p = 0.75). The potential difference in forward walking speed when far from the step likely derives from slower preferred speeds in darkness and not due to visual detection and planning for the step obstacle.

Our findings suggest that vision does not impact the ability of Argentine ant workers to identify and cross step obstacles. Ants often decelerated only when their antennae reached the step (Fig. 5A, S7); this behavior likely represents a reliance on tactile sensation instead of vision. Antennal-based strategies have been observed in tiger beetles (Zurek and Gilbert, 2014), stick insects (Schütz and Dürr, 2011), and cockroaches (Baba et al., 2010; Gart and Li, 2018; Harley et al., 2009), with all observed to adjust speed or body positioning before reaching obstacles. Our observations that Argentine ants decelerate to slower speeds, but do not stop as the mandible nears the step edge (Fig. 5A) differs from maximally sprinting cockroaches that collide with a tall vertical obstacle before climbing (Jayaram et al., 2018). Argentine ant workers appear to decelerate of their own control rather than through interactions with an obstacle.

Worker ants demonstrated the ability to rapidly decelerate once in the vicinity of a step. To generate these body accelerations, ants likely generate large friction forces through friction-inducing hairs on their distal tarsi and an actively-controlled adhesive organ next to their claws (Endlein and Federle, 2015). As a consequence, ants are likely capable of decelerating from their average speed (25.9 mm/s on flat ground) to rest within a single step. The latency between an antennal contact and reactionary behavior has not been measured in ants, however reaction times are ∼40 ms in stick insects (Schütz and Dürr, 2011), 25 ms in cockroaches (Ye and Comer, 1996; Ye et al., 2003), and <10 ms in hawk moths (Krishnan and Sane, 2014). Argentine ants walk at 5-15 strides per second, corresponding to a minimum stride duration of 66 ms (Clifton et al., 2020). Therefore it is likely that tactile sensation from the antennae is sufficiently fast to enable decelerations within one stride of encountering an obstacle.

Argentine ants demonstrate no difference in walking performance due to light conditions, however, most tested insects show some influence from vision. Grasshoppers slow down before even their antennae reach an obstacle (Pearson and Franklin, 1984) and miss targeted footholds with their forelimbs more frequently when blinded (Niven et al., 2010). In darkness, blow flies alter body posture and use their forelegs to probe surfaces, reducing walking speeds by 50% on rough terrain (Kress and Egelhaaf, 2012). Cockroaches with occluded ocelli prefer to tunnel under versus climb over an obstacle (Harley et al., 2009), while cockroaches with occluded compound eyes collide more frequently with obstacles than when their vision is unimpeded (Baba et al., 2010). To our knowledge, only two studies show no influence of vision: cockroaches sprinting (approximately 25 body length/second) then transitioning to vertical climbing (Jayaram et al., 2018) and a diurnal species of tiger beetle effectively “blinded” by motion blur due to fast running speeds (up to 120 body lengths/second) (Zurek and Gilbert, 2014). Unlike these two examples, Argentine ant workers walk at moderate speeds (approximately 10 body lengths/second) yet also maintain similar walking behavior between light and dark conditions. This observation could be explained by low resolution eyes in Argentine ants that preclude high-resolution vision even in light conditions. However, robotic tests show that even low resolution vision can permit successful navigation (Milford, 2013). Alternatively, Argentine ant workers could derive all necessary sensory feedback for walking from chemosensory and proprioceptive cues using the antennae and limbs.

### Antennal activity does not differ in light and darkness

In the absence of light, ants may still gather sensory information about substrate structure through tactile contact of the antennae and limbs. If Argentine ant workers use vision for walking, we expect that their antennal activity would increase in darkness to compensate for lost visual information. We found that on the uneven substrate, antennal speed with respect to the body was 35.1% faster than on flat ground (LME chi-squared test, p < 0.001) (Fig. 6B), however lighting did not have a significant effect (LME chi-squared test, p = 0.17). Counter to our expectations, antennal speed did not increase in darkness, with a 0.2% and 1.0% decrease on flat and uneven terrain respectively. We also found that antennal speeds increased for ants close to a step (LME chi-squared test, p < 0.001), but with no significant impact of step contrast (p = 0.03) or lighting (p = 0.84).

**Figure 6.**
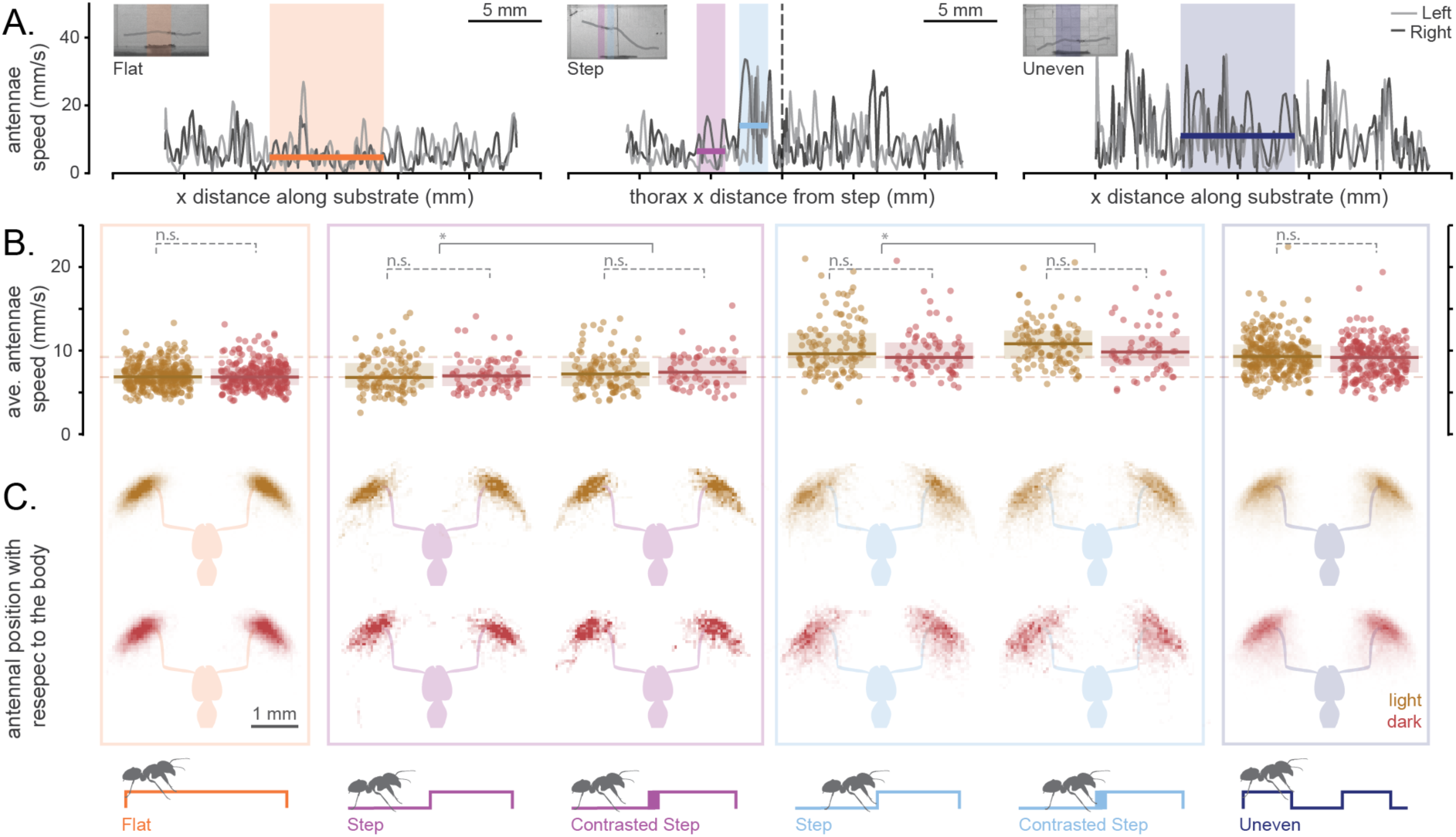
Antennal function for ants walking on flat or uneven terrain and while crossing a 1 mm step obstacle. (A) Antennal speeds with respect to the body of the ant were averaged across 8 mm on flat (orange) and uneven (navy) substrates and while ants were far (4-6 mm, purple) and close (1-3 mm, light blue) from the step. (B) Distributions of observed antennal speeds in light (yellow) and dark (red) conditions. Boxes represent the median and interquartile range. Dashed lines show the median antennal speed values for light (yellow) and dark (red) conditions on flat (lower) and uneven (upper) terrain. At far distances from the step, antennal speeds resemble those on flat ground. Close to the step, antennal speeds slightly exceed those observed on uneven ground. (C) Antennal spatial patterns appear similar under light and dark conditions, but cover a larger range when on uneven terrain or when close to the step. Opacity shows relative spatial density.

Qualitatively, Argentine ant workers moved their antennae over a larger area while walking on uneven terrain or while close to the step (Fig. 6C), however lighting conditions did not appear to influence exploration range. The observed spatial antennae patterns in Argentine ants on flat ground mimic carpenter ant “trail following” behavior, while spatial patterns during walking near a step obstacle and on uneven terrain resembled “exploration” (Draft et al., 2018). Our findings suggest that antennal function and ant behavior likely shift as a result of substrate unevenness, but are unaffected by vision in Argentine ants. Unlike observations in carpenter ants, we find that Argentine ants did not tightly correlate antennal motion relative to the head (ρ< 0.5, Fig. S3C-F). Since antennae evolved from modified appendages (Krishnan and Sane, 2015), variation in antennal coordination on uneven terrain may reveal neural control mechanisms associated with derived functions.

Apart from a very small decrease in antennal speed in darkness for Argentine ants walking uneven terrain, lighting condition did not influence overall antennal activity. Ant antennae contain hundreds of sensory receptors (Nakanishi et al., 2009), gathering critical tactile and chemosensory information about the environment. Ant antennae function in determining pheromone trail location (Draft et al., 2018), trail polarity (Jackson et al., 2004), nest location (Steck et al., 2011), and nestmate recognition (Frasnelli et al., 2012). In other insects, antennae have also been shown to facilitate wall following (Mongeau et al., 2013) and obstacle identification (Zurek and Gilbert, 2014). For cockroaches and stick insects, vision directs antennal motion towards obstacles (Comer and Baba, 2011) but has not been directly tested for an influence on the highly-coordinated antennal motions observed during walking (Dürr et al., 2001; Krause et al., 2013; Okada and Toh, 2004). Both sighted and blinded stick insects retarget their feet after sensing an obstacle with their antennae, suggesting that antennal sensation may be sufficient for navigating uneven terrain (Schütz and Dürr, 2011). Our findings for Argentine ants similarly support a dominant, albeit context-specific, role of antennal activity during walking.

## Conclusions

Despite possessing large eyes compared to closely-related species (Wild, 2004), Argentine ant workers do not appear to use vision while walking on flat and uneven terrain, or while crossing over step obstacles. In contrast, terrain unevenness and obstacles prompted large changes in walking speed and antennal activity, highlighting the importance of considering nontraditional laboratory conditions when studying movement control and behavior.

Our findings combined with results from a previous study testing Argentine ant navigation (Aron et al., 1993) indicate that workers do not use vision for walking or navigation. Given that eyes are costly to maintain (Niven et al., 2007) and given numerous independent examples of ant species with either highly reduced eyes or no eyes at all (Deharveng and Bedos, 2018; Pape and University of Arizona, 2016; Tierney et al., 2018), the presence of eyes in Argentine ants seems at odds with our findings. Vision perhaps could play a role in short-range perception or perceiving light patterns to entrain circadian rhythms. Alternatively, vision might play a crucial role for Argentine ant queens or males (which both have relatively large eyes), remaining in the worker caste due to developmental constraints.

We find that darkness did not compromise Argentine ant walking performance. As such, workers foraging at night can benefit from reduced risks of thermal stress or predation (Greiner, 2006; Wcislo et al., 2004). Argentine ant workers likely rely on tactile or chemical cues to inform walking strategies and obstacle negotiation. Antennae and proprioceptive limbs likely provide sensory feedback to ensure secure foot placement and limb coordination while maintaining relatively fast walking speeds (approximately 10 body lengths per second) (Zill et al., 2010). These strategies may provide relevant inspiration for legged robotics and autonomous vehicles (Cowan et al., 2005; Rudy et al., 2013). Furthermore, our findings demonstrate that possessing eyes does not necessarily correspond to an active role of vision in every task. This dissociation highlights the importance of testing functional interpretations for anatomical structures.

## Methods

### Linepithema humile

Argentine ants (Linepithema humile) were collected from six locations in San Diego county during February and March 2020. Argentine ants are broadly considered to form a super-colony in Southern California (Tsutsui et al., 2003), so our reference to “colonies” refers to varying collection date and location. Once a large aggregation of ants was found, 100-500 workers and surrounding soil were transferred to a plastic container and stored in a custom recording arena in the laboratory. Ants were housed overnight until recording began the following morning. Ants were housed in the lab for ∼30 hours total. Data from one collection day was removed from analysis since many ants congregated within the tunnel.

### Details of experiment and recording

Ant colonies were collected between 7:00 and 10:00 AM then allowed to acclimate for at least seven hours in the lab before opening a pathway through a 3D-printed tunnel to a foraging arena (Fig. 2A). The foraging arena contained food made from sugar, water, and polymer crystals (to delay evaporation). After opening the foraging arena, the set-up was not disturbed for the remainder of the experiment. Overnight, the ants explored the foraging arena and developed a recruitment trail through the tunnel. Each colony was then filmed during two recording sessions, from 7:00 AM - 2:00 PM on subsequent days. Temperature recordings were collected every 30 seconds while recording (Jepeak BE300263), ranging between 25.6875 and 25.9375 C. After the recording session, the colony was released back to its collection location and the tunnel and substrates were cleaned in warm, soapy water. The equipment dried overnight, dissipating any collected pheromone trails before the first recording session for the next collected colony.

For each colony, four substrates were randomly positioned in the tunnel leading from the colony container to the foraging arena. In addition to a flat substrate, we included two substrates with a single 1 mm step (one with the edge of the step painted black using a sharpie 12 hours before testing so it could dry) and an uneven checkerboard with a box width of 3 mm and step height of 1 mm. A step height of 1 mm approximates the hip height of Argentine ant workers (estimated from publicly-available photos of live ants), therefore challenging the ant without requiring vertical climbing.

During a recording session, two web cameras (YoLuke A860-Blue, Jide Technology Co., Ltd., China) focused on the tunnel automatically detected an incoming ant and triggered one of four machine vision cameras (Blackfly S 13Y3M, PointGrey, Inc., Canada). The cameras were each attached to a varifocal lens (20-100mm, 13VM20100AS, Tamron Co., Ltd., Japan). After being triggered by a web camera, each machine vision camera recorded for 3 seconds at 240 fps (720 frames total, 1000 x 500 pixels) then was paused for 80 seconds to reduce the probability of re-recordings the same ant. A still frame from the triggering web camera was saved in association with every video. In total, we recorded and analyzed over 4,200 videos.

The experimental set-up was illuminated by white LEDs (Lighting Ever, Daylight White, 6000K) on a 12:12 light:dark cycle. The tunnel was backlit using infrared LEDs (SMD3528-300-IR, 850 nm). During the recording session, the white LEDs were turned off every hour for 30 minutes using a wall outlet timer (15119, General Electric). During these dark periods, two IR flood lights (850 nm; Tendelux, Shenzhen, China) turned on to illuminate the tunnel from the side, enabling ant detection by the web cameras. Ants possess two or three photoreceptors (Ogawa et al., 2015). Although the spectral sensitivities of Argentine ants has not been measured, no ant species has demonstrated a sensitivity to light above 650 nm (Aksoy and Camlitepe, 2018; Briscoe and Chittka, 2001). Therefore, the use of 850 nm lighting most likely acts as complete darkness for Argentine ants. The illumination of the two lighting conditions were <1.8 vs. 200-2,500 lux (Extech LT40, NH, USA). The measured light lux levels ranged based on the orientation of the luxmeter, which was held at the tunnel location and rotated replicating the possible facings of the ants while walking. These values align with light levels measured for bull ants foraging on a dark evening and in bright light (Narendra et al., 2013b). A separate dataset of nine colonies was collected in August 2019 comparing dark and dusk light levels (60 lux), with a summary figure in the Supplemental Information (Figs. S4). Findings from the dusk light dataset aligned with those presented here, with the exception that ants walked on average at faster speeds. Speed variation might result from collecting colonies during the summer, due to the dramatic seasonal cycles within Argentine ant colonies (Markin, 1970).

To quantify the facet dimensions of Argentine ant workers, we painted the heads of 20 individuals with clear nail polish (as in (Narendra et al., 2016)). We removed the nail polish molds and visualized each eye using a microscope (Olympus BX51, 40x). The eye molds were not flat and unable to be flattened with release cuts due to their small size. Instead we took photos at multiple focal planes (> 10 per eye) and focus stacked the images (Python, https://github.com/cmcguinness/). We selected the 5 clearest eyes for analysis. The facets in each eye were manually outlined and measured in ImageJ (Schneider et al., 2012), using a calibration from photos of a resolution test target (Thorlabs Inc., NJ, U.S.A., NBS 1952).

### Overview of tracking ants in highspeed videos

Each video was analyzed to track the body and antennae of all ants in view. This consisted of three steps. First, ant-centered videos were generated by grossly estimating ant locations in each frame then associating identified ants across frames, resulting in “trackways”. Second, trackways were used to generate ant-centered videos. Third, the ant-centered videos were processed using a deep-learning based tracking software (Pereira et al., 2019), outputting the locations of 4 landmarks along the body and three points along the antennae. These data were then processed to remove any likely poor tracking by the program. Each step is detailed below.

### Full-body tracking

Tracking ant locations in each frame was achieved by identifying all ants in each frame and associating individuals across frames. Briefly, to identify ants in each frame, we (1) normalized the video using background division, (2) used image processing to isolate the body of each ant, and (3) fit contours to each ant body to estimate the location and orientation. Ants identified in each frame were associated across frames using a Kalman filter generating “trackways” (Straw et al., 2011). For details see the supplementary materials in (Clifton et al., 2020).

### Generation of ant-centered videos

The facing of the ant in each kalman-associated “trackway” was estimated by attempting to find the asymmetries in the isolated contour due to the antennae. Then ant orientation was processed to remove any 90° or 180° jumps due to errors. Once the orientation was consistent throughout a trackway, it was filtered using a moving average (window size = 11 frames, ignoring any windows with fewer than 2 non-nan values). The x- and y-coordinates of the trackways were also filtered using a low-pass butterworth filter (Scipy, n=2, □_n_ = 0.2).

Each trackway was now used to generate ant-centered images with a dark background. Each background-divided frame was rotated so that the direction of the ant facing aligned with the +x direction (imutils.rotate_bound, Python) and cropped to a shape of 200×200 pixels around the center of the ant. Each cropped, background-divided image was then inverted and adjusted to increase the contrast between ant and background. To remove any other ants in the cropped frame, we identified any non-central, large foreground objects, and removed those which intersected with the edge of the cropped frame. Further details can be found in (Clifton et al., 2020). This process resulted in a video with an isolated, light-colored ant centered in the frame but walking towards the right against a dark background.

While our method of finding the head-orientation of the ant in each frame was mostly accurate, occasionally we generated video with the ant facing towards the left. To identify and flip these videos, we used a support vector machine (SVM). A training group of 1345 ant-centered pictures were used to train the SVM to classify 3 ant facing categories: right (ant faces in +x direction), left (ant faces in -x direction), or blank (the ant-centered picture is completely black as would be generated if the ant facing angle = nan). The first 100 frames of each trackway cropped video were classified as left, right, or blank. If the number of left-facing images outnumbered the right-facing images, the ant facing angles and ant-centered images for the trackway were rotated by 180°.

### “LEAP” deep-learning tracking of body and antennae

We used a recent deep-learning approach, implemented in MATLAB and Python (Pereira et al., 2019), to track landmarks on the ant in each video. We specified a skeleton of connected points to track, including six along the body (gaster tip, waist, neck, mandible, eyes) and three along each antennae. We manually identified these points in a training set of 140 frames. The LEAP tracker then predicted the landmark locations in all videos, requiring between 3 and 30 seconds depending on the number of frames. The LEAP network was trained using the following settings: scale = 1, kernel for confidence maps = 5, mirrored images enabled, leap_cnn network architecture, 64 base filters, 5 deg rotation angle, 25 epochs, 50 batches per epoch, 50 samples per batch, validation fraction = 0.1, AMSGRAD enabled, learning rate reduction factor = 0.1 after 3 epochs with the learning rate changing by less than 1e-5.

The general kinematic variability of ant walking, especially on uneven terrain, resulted in some inaccurate tracking predictions. Identifying and removing these points was a multistep process, with full details in the supplementary methods of (Clifton et al., 2020). Briefly, predicted landmarks with a low confidence value (output from the LEAP tracker) were removed along with points that jumped by more than 10 pixels (∼0.33mm) within one frame. We also removed any datapoint that deviated from the average position of that landmark (calculated using the middle 50% of points) by more than ⅔ of the middle 50% range. The x- and y-coordinates of each landmark were then low pass filtered (Scipy, butterworth, n=2, □_n_ = 0.3), while removing sections of the tracking with fewer than 9 consecutive non-nan values. If after this processing, fewer than 50 finite (non-nan) data points remained in the full landmark trace, that trace was removed from further analysis. Past comparisons of this approach with manually-tracked landmarks, demonstrated a high accuracy of the resulting post-processed data points.

### General analysis

The lighting condition for each video was determined by evaluating the associated web camera still frames. The lighting timer was not accurate in switching the lighting exactly every 30 minutes, so instead the timing of the lighting switches were determined by comparing the average illumination in each photo. Any videos within 5 seconds of a lighting switch were removed from further analysis.

Ants close to the edge of the video frame may be partially off-screen, therefore with unreliable tracking. Similarly, ants near the edge of a substrate could behave usually due to the transition from the flat tunnel floor. To identify these instances, image processing was used to locate the dark edges of the substrate and the step in each video. Any data points where the center of the ant was within 2 mm (∼64 pixels) of a substrate edge were removed. The information on the step-location was later used for analyzing ant walking while approaching and climbing the step.

While ants primarily walked through the tunnel to and from the foraging arena, they displayed some behavioral variation, including antennal cleaning and conspecific interactions. To identify and be able to remove these stationary behaviors from future analysis (e.g. walking speed comparisons), we low-pass filtered the instantaneous speed (Scipy, butterworth, n=2, □_n_ = 0.6) and removed any sections where this average walking speed dipped below 3 mm/s.

### Body speed and trajectory analysis, flat and uneven substrates

The low-pass filtered x- and y-coordinates of the waist throughout each trackway (excluding slow behaviors as described above) were used to calculate instantaneous velocity, which was then low-pass filtered again to remove the large accelerations present in ant walking (see Fig. 3B). The resulting velocity was decomposed relative to the orientation of the ant, generating a forward speed. The distance traveled at each speed was used to create a “histogram” of walking speeds (Fig. 1C). The preferred, “average” speed was calculated from the median of the distribution, while the top speed was determined from the cut-off for the fastest 5% of the total distance traveled under each condition (e.g. light on flat substrate). To test if the distributions differed from each other, we used a Mann-Whitney rank test, with 2 mm of distance traveled representing one data-point.

To test if the trajectories of the ants differed on flat vs. uneven substrates or under light vs. dark conditions, we calculated the straightness, sinuosity, and fractal dimension for each trackway.

- Straightness was defined as the total distance traveled divided by the net distance traveled.
- Sinuosity was calculated based on (Bovet and Benhamou, 1988). Briefly, each trackway section was split into approximately 2 mm steps (based off of an average 2 mm step length observed in (Clifton et al., 2020). The turning angle traveled over each step (φ) was used to determine the mean cosine (*c*) and mean sine (*s*) of the changes of direction. From there sinuosity was calculated as:

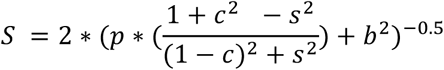

where *p* is the average step length (2 mm) and *b* is the standard deviation of step length errors compared to the average step length. Finding *b* resulted from how we identified the ant location at each step. The pathway was divided into approximately 2 mm steps by identifying when the actual distance traveled approached a multiple of 2 mm. The errors of the actual step lengths from an ideal 2 mm were compiled and the standard deviation of this distribution represented *b*. We also re-calculated sinuosity using *b* = 0.5 mm, and saw no change in the results.
- The fractal dimension of each trackway section was determined by segmenting the trackway by multiple lengths and comparing the net distances traveled for those segments to the segment lengths. For example a 7 mm long trackway was segmented into 0.15 mm travel distances and the net distances traveled for all sections were summed. This was repeated for sections of 0.63, 1.10, 1.58, 2.05, 2.52, 2.99, and 3.47 mm. A linear regression (Scipy, linregress) of the summed net distances vs. section length resulted in a slope of 2.07. The fractal dimension (*D*) was defined as *D =* 1 - slope. The segment lengths chosen for each trackway were chosen as ranging inclusively from 0.15 mm to half of the total distance traveled, with a step value of the total distance traveled/15 (i.e. [0.15 to d_total/2 by d_total/15]).

These values were calculated for all sections of each ant trackway then averaged. Any sections with fewer than 20 data points (< 83.3ms) or where the ant traveled < 3 mm were excluded. For sinuosity and fractal dimension calculations, any time there wasn’t a location of the ant within 2 pixels (0.06 mm) of the segmented distance (e.g. every 2 mm for sinuosity), the trackway section was excluded.

To determine if there was a significant influence of lighting (light vs. dark) on tackway tortuosity, we used nonparametric rank testing. The distributions of straightness, sinuosity, and fractal dimensions were highly skewed and unable to be normalized for parametric testing. Instead, the data for each colony and substrate (flat vs. array) were analyzed using a Mann-Whitney U test, testing the factor of light (wilcox.test(Y∼light), R). The resulting p-values are listed in Table S2.

### Step crossing analysis

To determine if step crossing performance shifts under light and dark conditions, we calculated four measures. We only included trials where the ant stepped upwards.

1. Step crossing time. We defined crossing as starting with the waist at a distance of 4 mm from the step (before any observed deceleration) and ending when the waist passes 1 mm away from the step.
2. Step crossing angle. We measured the net angular displacement of the ant from crossing starting to stopping. If an ant were to detect the step and veer to walk along the edge, this would be represented as a large crossing angle.
3. Forward speed far from step. The instantaneous forward speed of each ant was low-pass filtered (Scipy, butterworth, n=2, □_n_ = 0.6). We calculated the median low-pass filtered speed of each ant when the waist was between 6 and 4 mm before the step. Any trials with fewer than 10 data points within this window were removed.
4. Forward speed close to step. We calculated the median low-pass filtered speed of each ant when the waist was between 3 and 1 mm before the step. Any trials with fewer than 10 data points within this window were removed.

To statistically test for differences in these parameters due to substrate type (contrasted vs. uncontrasted steps) and lighting (light vs. dark), we used linear mixed-effect modeling (lme4::lmer function in R)(Bates et al., 2015). Crossing time and the speed close to the step were normalized using a logarithmic function. The goodness of fit of these models was confirmed by examining quantile-quantile and residual plots. We used the following full models:

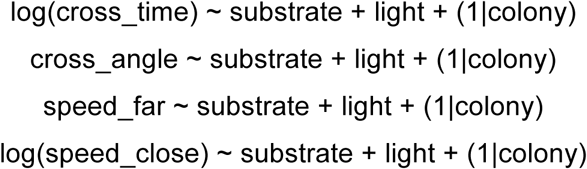

These models were compared to reduced models, removing either the substrate or light factors, and analyzed with a Chi-squared Likelihood Ratio test (anova function in R).

### Antennal activity analysis

Average antennal speeds were calculated on flat and uneven substrates, and with ants far and close to step obstacles. For flat and uneven terrain, ants were analyzed as they walked through the center of the substrate, with the waist between 11 and 19 mm from the left edge of the 30 mm long substrate. The analysis windows on step substrates coincided with those used for average body speed far and close to the step, from 6-to-4 and 3-to-1 mm before the step respectively. The tracking of each antenna was referenced to the body of the ant using the vector from the waist to the neck. Instantaneous speed with respect to the body was used to find the median speed for each antenna, which were then averaged.

The average antennal speed on flat and uneven terrain were analyzed for an influence of light and substrate in a similar manner to the step crossing metrics above. The exact linear mixed-effect models were:

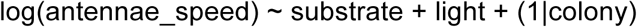

The model for step substrates included close vs. far distance as an additional variable:

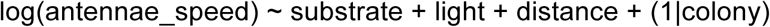

As above, Chi-squared Likelihood Ratio tests (anova function in R) tested the influence of each fixed variable.

### Data and code availability

The datasets generated during and analyzed during the current study are publicly available on Dryad at <private while under review>. Code used for analysis is available at https://github.com/gtclifton.

## Acknowledgments

We thank Prof. Michael Tolley for 3D printer access and Ben Shih and Chris Cassidy for printing help.

## Author contributions

All authors contributed to the conceptualization and methodology design for this study. G.C. performed data collection, analysis, and drafted the manuscript. All authors contributed to critically revising the manuscript. All authors gave final approval for publication and agree to be held accountable for the work performed therein.

## References

Abril, S., Oliveras, J. and Gómez, C. (2007). Foraging activity and dietary spectrum of the Argentine ant (Hymenoptera: Formicidae) in invaded natural areas of the northeast Iberian Peninsula. Environ. Entomol. 36, 1166–1173.

Aksoy, V. and Camlitepe, Y. (2018). Spectral sensitivities of ants – a review. Animal Biology 68, 55–73.

Almeida, P. J. A. L., Paulo J A, Vieira, M. V., Kajin, M., Forero-Medina, G. and Cerqueira, R. (2010). Indices of movement behaviour: conceptual background, effects of scale and location errors. Zoologia (Curitiba) 27, 674–680.

Aron, S., Beckers, R., Deneubourg, J. L. and Pasteels, J. M. (1993). Memory and chemical communication in the orientation of two mass-recruiting ant species. Insectes Sociaux 40, 369–380.

Baba, Y., Tsukada, A. and Comer, C. M. (2010). Collision avoidance by running insects: antennal guidance in cockroaches. J. Exp. Biol. 213, 2294–2302.

Baker, G. T. and Ma, P. W. K. (2006). Morphology and number of Ommatidia in the compound eyes of Solenopsis invicta, Solenopsis richteri, and their hybrid (Hymenoptera: Formicidae). Zoologischer Anzeiger - A Journal of Comparative Zoology 245, 121–125.

Bates, D., Mächler, M., Bolker, B. and Walker, S. (2015). Fitting Linear Mixed-Effects Models Using lme4. Journal of Statistical Software 67,.

Bender, J. A., Simpson, E. M., Tietz, B. R., Daltorio, K. A., Quinn, R. D. and Ritzmann, R. E. (2011). Kinematic and behavioral evidence for a distinction between trotting and ambling gaits in the cockroach Blaberus discoidalis. J. Exp. Biol. 214, 2057–2064.

Benhamou, S. (2004). How to reliably estimate the tortuosity of an animal’s path: straightness, sinuosity, or fractal dimension? J. Theor. Biol. 229, 209–220.

Bernstein, S. and Finn, C. (1971). Ant compound eye: Size-related ommatidium differences within a single wood ant nest. Experientia 27, 708–710.

Bingman, V. P., Graving, J. M., Hebets, E. A. and Wiegmann, D. D. (2017). Importance of the antenniform legs, but not vision, for homing by the neotropical whip spider. J. Exp. Biol. 220, 885–890.

Birn-Jeffery, A. V., Hubicki, C. M., Blum, Y., Renjewski, D., Hurst, J. W. and Daley, M. A. (2014). Don’t break a leg: running birds from quail to ostrich prioritise leg safety and economy on uneven terrain. J. Exp. Biol. 217, 3786–3796.

Blaesing, B. (2004). Stick insect locomotion in a complex environment: climbing over large gaps. Journal of Experimental Biology 207, 1273–1286.

Bovet, P. and Benhamou, S. (1988). Spatial analysis of animals’ movements using a correlated random walk model. Journal of Theoretical Biology 131, 419–433.

Briscoe, A. D. and Chittka, L. (2001). The evolution of color vision in insects. Annu. Rev. Entomol. 46, 471–510.

Cheng, K., Narendra, A. and Wehner, R. (2006). Navigation in the Central Australian Desert Ant Melophorus bagoti: Some Initial Results. PsycEXTRA Dataset.

Clifton, G. T., Holway, D. and Gravish, N. (2020). Uneven substrates constrain walking speed in ants through modulation of stride frequency more than stride length. Royal Society Open Science 7, 192068.

Collett, T. S. (2002). Insect vision: controlling actions through optic flow. Curr. Biol. 12, R615–7.

Comer, C. and Baba, Y. (2011). Active touch in orthopteroid insects: behaviours, multisensory substrates and evolution. Philos. Trans. R. Soc. Lond. B Biol. Sci. 366, 3006–3015.

Cowan, N. J., Ma, E. J., Cutkosky, M. and Full, R. J. (2005). A Biologically Inspired Passive Antenna for Steering Control of a Running Robot. Springer Tracts in Advanced Robotics 541–550.

Creamer, M. S., Mano, O. and Clark, D. A. (2018). Visual Control of Walking Speed in Drosophila. Neuron 100, 1460–1473.e6.

Cronin, T. W., Johnsen, S., Justin Marshall, N. and Warrant, E. J. (2014). Visual Ecology.

Deharveng, L. and Bedos, A. (2018). Diversity of Terrestrial Invertebrates in Subterranean Habitats. Cave Ecology 107–172.

Draft, R. W., McGill, M. R., Kapoor, V. and Murthy, V. N. (2018). Carpenter ants use diverse antennae sampling strategies to track odor trails. J. Exp. Biol. 221,.

Dürr, V. (2001). Stereotypic leg searching movements in the stick insect: kinematic analysis, behavioural context and simulation. J. Exp. Biol. 204, 1589–1604.

Dürr, V., König, Y. and Kittmann, R. (2001). The antennal motor system of the stick insect Carausius morosus: anatomy and antennal movement pattern during walking. J. Comp. Physiol. A 187, 131–144.

Endlein, T. and Federle, W. (2015). On Heels and Toes: How Ants Climb with Adhesive Pads and Tarsal Friction Hair Arrays. PLoS One 10, e0141269.

Fewell, J. H. (1988). Energetic and time costs of foraging in harvester ants, Pogonomyrmex occidentalis. Behav. Ecol. Sociobiol. 22, 401–408.

Frasnelli, E., Iakovlev, I. and Reznikova, Z. (2012). Asymmetry in antennal contacts during trophallaxis in ants. Behav. Brain Res. 232, 7–12.

Frolov, R. V. and Ignatova, I. I. (2020). Electrophysiological adaptations of insect photoreceptors and their elementary responses to diurnal and nocturnal lifestyles. J. Comp. Physiol. A Neuroethol. Sens. Neural Behav. Physiol. 206, 55–69.

Frolov, R. V., Matsushita, A. and Arikawa, K. (2017). Not flying blind: a comparative study of photoreceptor function in flying and non-flying cockroaches. J. Exp. Biol. 220, 2335–2344.

Full, R. J. and Koditschek, D. E. (1999). Templates and anchors: neuromechanical hypotheses of legged locomotion on land. J. Exp. Biol. 202, 3325–3332.

Full, R. J. and Tu, M. S. (1990). Mechanics of six-legged runners. J. Exp. Biol. 148, 129–146.

Gart, S. W. and Li, C. (2018). Body-terrain interaction affects large bump traversal of insects and legged robots. Bioinspir. Biomim. 13, 026005.

Gilbert, C. (1997). Visual control of cursorial prey pursuit by tiger beetles (Cicindelidae). Journal of Comparative Physiology A: Sensory, Neural, and Behavioral Physiology 181, 217–230.

Graham, D. (1972). A behavioural analysis of the temporal organisation of walking movements in the 1st instar and adult stick insect (Carausius morosus). Journal of Comparative Physiology 81, 23–52.

Graham, P. and Cheng, K. (2009). Which portion of the natural panorama is used for view-based navigation in the Australian desert ant? J. Comp. Physiol. A Neuroethol. Sens. Neural Behav. Physiol. 195, 681–689.

Graham, P. and Philippides, A. (2017). Vision for navigation: What can we learn from ants? Arthropod Struct. Dev. 46, 718–722.

Gravish, N., Monaenkova, D., Goodisman, M. A. D. and Goldman, D. I. (2013). Climbing, falling, and jamming during ant locomotion in confined environments. Proc. Natl. Acad. Sci. U. S. A. 110, 9746–9751.

Greiner, B. (2006). Adaptations for nocturnal vision in insect apposition eyes. Int. Rev. Cytol. 250, 1–46.

Greiner, B., Narendra, A., Reid, S. F., Dacke, M., Ribi, W. A. and Zeil, J. (2007). Eye structure correlates with distinct foraging-bout timing in primitive ants. Curr. Biol. 17, R879–80.

Gronenberg, W. and Hölldobler, B. (1999). Morphologic representation of visual and antennal information in the ant brain. J. Comp. Neurol. 412, 229–240.

Harley, C. M., English, B. A. and Ritzmann, R. E. (2009). Characterization of obstacle negotiation behaviors in the cockroach, Blaberus discoidalis. J. Exp. Biol. 212, 1463–1476.

Haselsteiner, A. F., Gilbert, C. and Wang, Z. J. (2014). Tiger beetles pursue prey using a proportional control law with a delay of one half-stride. J. R. Soc. Interface 11, 20140216.

Hebets, E. A. (2002). Relating the unique sensory system of amblypygids to the ecology and behavior of Phrynus parvulus from Costa Rica (Arachnida, Amblypygi). Canadian Journal of Zoology 80, 286–295.

Heglund, N. C., Taylor, C. R. and McMahon, T. A. (1974). Scaling stride frequency and gait to animal size: mice to horses. Science 186, 1112–1113.

Heimonen, K., Immonen, E.-V., Frolov, R. V., Salmela, I., Juusola, M., Vähäsöyrinki, M. and Weckström, M. (2012). Signal coding in cockroach photoreceptors is tuned to dim environments. J. Neurophysiol. 108, 2641–2652.

Hogg, B. N., Nelson, E. H., Hagler, J. R. and Daane, K. M. (2018). Foraging Distance of the Argentine Ant in California Vineyards. J. Econ. Entomol. 111, 672–679.

Höltje, M. and Hustert, R. (2003). Rapid mechano-sensory pathways code leg impact and elicit very rapid reflexes in insects. J. Exp. Biol. 206, 2715–2724.

Howard, C. E., Chen, C.-L., Tabachnik, T., Hormigo, R., Ramdya, P. and Mann, R. S. (2019). Serotonergic Modulation of Walking in Drosophila. Curr. Biol. 29, 4218–4230.e8.

Human, K. G. and Gordon, D. M. (1996). Exploitation and interference competition between the invasive Argentine ant, Linepithema humile, and native ant species. Oecologia 105, 405–412.

Hurlbert, A. H., Ballantyne, F. and Powell, S. (2008). Shaking a leg and hot to trot: the effects of body size and temperature on running speed in ants. Ecological Entomology 33, 144–154.

Ignatova, I. I., Saari, P. and Frolov, R. V. (2020). Latency of phototransduction limits transfer of higher-frequency signals in cockroach photoreceptors. J. Neurophysiol. 123, 120–133.

Isakov, A., Buchanan, S. M., Sullivan, B., Ramachandran, A., Chapman, J. K. S., Lu, E. S., Mahadevan, L. and de Bivort, B. (2016). Recovery of locomotion after injury in Drosophila melanogaster depends on proprioception. J. Exp. Biol. 219, 1760–1771.

Jackson, D. E., Holcombe, M. and Ratnieks, F. L. W. (2004). Trail geometry gives polarity to ant foraging networks. Nature 432, 907–909.

Jayaram, K., Mongeau, J.-M., Mohapatra, A., Birkmeyer, P., Fearing, R. S. and Full, R. J. (2018). Transition by head-on collision: mechanically mediated manoeuvres in cockroaches and small robots. J. R. Soc. Interface 15,.

Jayatilaka, P., Narendra, A., Reid, S. F., Cooper, P. and Zeil, J. (2011). Different effects of temperature on foraging activity schedules in sympatric Myrmecia ants. J. Exp. Biol. 214, 2730–2738.

Kaspari, M. and Weiser, M. D. (1999). The size–grain hypothesis and interspecific scaling in ants. Functional Ecology 13, 530–538.

Klotz, J. H. and Reid, B. L. (1992). The use of spatial cues for structural guideline orientation inTapinoma sessile andCamponotus pennsylvanicus (Hymenoptera: Formicidae). Journal of Insect Behavior 5, 71–82.

Klotz, J. H., Reid, B. L. and Gordon, W. C. (1992). Variation of ommatidia number as a function of worker size inCamponotus pennsylvanicus (DeGeer) (Hymenoptera: Formicidae). Insectes Sociaux 39, 233–236.

Knaden, M. and Graham, P. (2016). The Sensory Ecology of Ant Navigation: From Natural Environments to Neural Mechanisms. Annu. Rev. Entomol. 61, 63–76.

Krause, A. F., Winkler, A. and Dürr, V. (2013). Central drive and proprioceptive control of antennal movements in the walking stick insect. J. Physiol. Paris 107, 116–129.

Kress, D. and Egelhaaf, M. (2012). Head and body stabilization in blowflies walking on differently structured substrates. J. Exp. Biol. 215, 1523–1532.

Krishnan, A. and Sane, S. P. (2014). Visual feedback influences antennal positioning in flying hawk moths. J. Exp. Biol. 217, 908–917.

Krishnan, A. and Sane, S. P. (2015). Antennal Mechanosensors and Their Evolutionary Antecedents. Advances in Insect Physiology 59–99.

Land, M. F. (1997a). VISUAL ACUITY IN INSECTS. Annual Review of Entomology 42, 147–177.

Land, M. F. (1997b). THE RESOLUTION OF INSECT COMPOUND EYES. Israel Journal of Plant Sciences 45, 79–91.

Land, M. F. and Collett, T. S. (1974). Chasing behaviour of houseflies (Fannia canicularis). Journal of Comparative Physiology 89, 331–357.

Lee, T., Jang, S., Jeong, M. and Cho, D.-I. D. (2016). Allometric scaling of insects and animals for biomimetic robot design considerations. 2016 16th International Conference on Control, Automation and Systems (ICCAS).

Logan, D., Kiemel, T., Dominici, N., Cappellini, G., Ivanenko, Y., Lacquaniti, F. and Jeka, J. J. (2010). The many roles of vision during walking. Exp. Brain Res. 206, 337–350.

Markin, G. P. (1970). The Seasonal Life Cycle of the Argentine Ant, Iridomyrmex humilis (Hymenoptera: Formicidae), in Southern California. Annals of the Entomological Society of America 63, 1238–1242.

Menzi, U. (1987). Visual adaptation in nocturnal and diurnal ants. Journal of Comparative Physiology A 160, 11–21.

Milford, M. (2013). Vision-based place recognition: how low can you go? The International Journal of Robotics Research 32, 766–789.

Mongeau, J.-M., Demir, A., Lee, J., Cowan, N. J. and Full, R. J. (2013). Locomotion- and mechanics-mediated tactile sensing: antenna reconfiguration simplifies control during high-speed navigation in cockroaches. J. Exp. Biol. 216, 4530–4541.

Nakanishi, A., Nishino, H., Watanabe, H., Yokohari, F. and Nishikawa, M. (2009). Sex-specific antennal sensory system in the ant Camponotus japonicus: structure and distribution of sensilla on the flagellum. Cell Tissue Res. 338, 79–97.

Nams, V. O. (1996). The VFractal: a new estimator for fractal dimension of animal movement paths. Landscape Ecology 11, 289–297.

Narendra, A., Raderschall, C. A. and Robson, S. K. A. (2013a). Homing abilities of the Australian intertidal ant Polyrhachis sokolova. Journal of Experimental Biology 216, 3674–3681.

Narendra, A., Reid, S. F. and Raderschall, C. A. (2013b). Navigational efficiency of nocturnal Myrmecia ants suffers at low light levels. PLoS One 8, e58801.

Narendra, A., Ramirez-Esquivel, F. and Ribi, W. A. (2016). Compound eye and ocellar structure for walking and flying modes of locomotion in the Australian ant, Camponotus consobrinus. Scientific Reports 6,.

Niven, J. E., Anderson, J. C. and Laughlin, S. B. (2007). Fly photoreceptors demonstrate energy-information trade-offs in neural coding. PLoS Biol. 5, e116.

Niven, J. E., Buckingham, C. J., Lumley, S., Cuttle, M. F. and Laughlin, S. B. (2010). Visual Targeting of Forelimbs in Ladder-Walking Locusts. Current Biology 20, 86–91.

Niven, J. E., Ott, S. R. and Rogers, S. M. (2012). Visually targeted reaching in horse-head grasshoppers. Proc. Biol. Sci. 279, 3697–3705.

Ogawa, Y., Falkowski, M., Narendra, A., Zeil, J. and Hemmi, J. M. (2015). Three spectrally distinct photoreceptors in diurnal and nocturnal Australian ants. Proc. Biol. Sci. 282, 20150673.

Okada, J. and Toh, Y. (2004). Spatio-temporal patterns of antennal movements in the searching cockroach. J. Exp. Biol. 207, 3693–3706.

Okada, J. and Toh, Y. (2006). Active tactile sensing for localization of objects by the cockroach antenna. J. Comp. Physiol. A Neuroethol. Sens. Neural Behav. Physiol. 192, 715–726.

Ozaki, M., Wada-Katsumata, A., Fujikawa, K., Iwasaki, M., Yokohari, F., Satoji, Y., Nisimura, T. and Yamaoka, R. (2005). Ant nestmate and non-nestmate discrimination by a chemosensory sensillum. Science 309, 311–314.

Palavalli-Nettimi, R. and Narendra, A. (2018). Miniaturisation decreases visual navigational competence in ants. J. Exp. Biol. 221,.

Pape, R. and University of Arizona (2016). The importance of ants in cave ecology, with new records and behavioral observations of ants in Arizona caves. International Journal of Speleology 45, 185–205.

Patla, A. E. and Greig, M. (2006). Any way you look at it, successful obstacle negotiation needs visually guided on-line foot placement regulation during the approach phase. Neurosci. Lett. 397, 110–114.

Pearson, K. G. and Franklin, R. (1984). Characteristics of Leg Movements and Patterns of Coordination in Locusts Walking on Rough Terrain. The International Journal of Robotics Research 3, 101–112.

Pearson, K. G., Stein, R. B. and Malhotra, S. K. (1970). Properties of action potentials from insect motor nerve fibres. J. Exp. Biol. 53, 299–316.

Pereira, T. D., Aldarondo, D. E., Willmore, L., Kislin, M., Wang, S. S.-H., Murthy, M. and Shaevitz, J. W. (2019). Fast animal pose estimation using deep neural networks. Nat. Methods 16, 117–125.

Pick, S. and Strauss, R. (2005). Goal-driven behavioral adaptations in gap-climbing Drosophila. Curr. Biol. 15, 1473–1478.

Pratt, S. C., Brooks, S. E. and Franks, N. R. (2001). The Use of Edges in Visual Navigation by the Ant Leptothorax albipennis. Ethology 107, 1125–1136.

Ramirez-Esquivel, F., Leitner, N. E., Zeil, J. and Narendra, A. (2017). The sensory arrays of the ant, Temnothorax rugatulus. Arthropod Struct. Dev. 46, 552–563.

Reinhardt, L. and Blickhan, R. (2014). Level locomotion in wood ants: evidence for grounded running. J. Exp. Biol. 217, 2358–2370.

Reynolds, R. F. and Day, B. L. (2005a). Visual guidance of the human foot during a step. The Journal of Physiology 569, 677–684.

Reynolds, R. F. and Day, B. L. (2005b). Rapid visuo-motor processes drive the leg regardless of balance constraints. Curr. Biol. 15, R48–9.

Rudy, R., Cohen, A. J., Pulskamp, J. S., Polcawich, R. G. and Oldham, K. R. (2013). Antenna-Like Tactile Sensor for Thin-Film Piezoelectric Micro-Robots. Volume 1: 15th International Conference on Advanced Vehicle Technologies; 10th International Conference on Design Education; 7th International Conference on Micro- and Nanosystems.

Schilman, P. E., Lighton, J. R. B. and Holway, D. A. (2005). Respiratory and cuticular water loss in insects with continuous gas exchange: comparison across five ant species. J. Insect Physiol. 51, 1295–1305.

Schneider, C. A., Rasband, W. S. and Eliceiri, K. W. (2012). NIH Image to ImageJ: 25 years of image analysis. Nat. Methods 9, 671–675.

Schütz, C. and Dürr, V. (2011). Active tactile exploration for adaptive locomotion in the stick insect. Philosophical Transactions of the Royal Society B: Biological Sciences 366, 2996–3005.

Schwarz, S., Narendra, A. and Zeil, J. (2011). The properties of the visual system in the Australian desert ant Melophorus bagoti. Arthropod Structure & Development 40, 128–134.

Sheehan, Z. B. V., Kamhi, J. F., Seid, M. A. and Narendra, A. (2019). Differential investment in brain regions for a diurnal and nocturnal lifestyle in Australian Myrmecia ants. J. Comp. Neurol. 527, 1261–1277.

Sherman, A. and Dickinson, M. H. (2004). Summation of visual and mechanosensory feedback in Drosophila flight control. J. Exp. Biol. 207, 133–142.

Smid, K. A. and den Otter, A. R. (2013). Why you need to look where you step for precise foot placement: the effects of gaze eccentricity on stepping errors. Gait Posture 38, 242–246.

Steck, K., Hansson, B. S. and Knaden, M. (2011). Desert ants benefit from combining visual and olfactory landmarks. J. Exp. Biol. 214, 1307–1312.

Straw, A. D., Branson, K., Neumann, T. R. and Dickinson, M. H. (2011). Multi-camera real-time three-dimensional tracking of multiple flying animals. J. R. Soc. Interface 8, 395–409.

Stürzl, W., Grixa, I., Mair, E., Narendra, A. and Zeil, J. (2015). Three-dimensional models of natural environments and the mapping of navigational information. J. Comp. Physiol. A Neuroethol. Sens. Neural Behav. Physiol. 201, 563–584.

Szczecinski, N. S., Büschges, A. and Bockemühl, T. (2018). Direction-Specific Footpaths Can Be Predicted by the Motion of a Single Point on the Body of the Fruit Fly Drosophila Melanogaster. Biomimetic and Biohybrid Systems 477–489.

Theunissen, L. M., Bekemeier, H. H. and Dürr, V. (2015). Comparative whole-body kinematics of closely related insect species with different body morphology. J. Exp. Biol. 218, 340–352.

Tierney, S. M., Langille, B., Humphreys, W. F., Austin, A. D. and Cooper, S. J. B. (2018). Massive Parallel Regression: A Précis of Genetic Mechanisms for Vision Loss in Diving Beetles. Integr. Comp. Biol. 58, 465–479.

Tsutsui, N. D., Suarez, A. V. and Grosberg, R. K. (2003). Genetic diversity, asymmetrical aggression, and recognition in a widespread invasive species. Proceedings of the National Academy of Sciences 100, 1078–1083.

van Zweden, J. S. and d’Ettorre, P. Nestmate recognition in social insects and the role of hydrocarbons. Insect Hydrocarbons 222–243.

Vega, S. Y. and Rust, M. K. (2003). Determining the Foraging Range and Origin of Resurgence After Treatment of Argentine Ant (Hymenoptera: Formicidae) in Urban Areas. Journal of Economic Entomology 96, 844–849.

Warrant, E. J. (2017). The remarkable visual capacities of nocturnal insects: vision at the limits with small eyes and tiny brains. Philos. Trans. R. Soc. Lond. B Biol. Sci. 372,.

Warzecha, A. and Egelhaaf, M. (2000). Response latency of a motion-sensitive neuron in the fly visual system: dependence on stimulus parameters and physiological conditions. Vision Res. 40, 2973–2983.

Watson, J., Ritzmann, R., Zill, S. and Pollack, A. (2002). Control of obstacle climbing in the cockroach, Blaberus discoidalis. I. Kinematics. Journal of Comparative Physiology A: Sensory, Neural, and Behavioral Physiology 188, 39–53.

Wcislo, W. T., Arneson, L., Roesch, K., Gonzalez, V., Smith, A. and Fernández, H. (2004). The evolution of nocturnal behaviour in sweat bees, Megalopta genalis and M. ecuadoria (Hymenoptera: Halictidae): an escape from competitors and enemies? Biological Journal of the Linnean Society 83, 377–387.

Wehner, R. and Muller, M. (2006). The significance of direct sunlight and polarized skylight in the ant’s celestial system of navigation. Proceedings of the National Academy of Sciences 103, 12575–12579.

Weihmann, T. and Blickhan, R. (2009). Comparing inclined locomotion in a ground-living and a climbing ant species: sagittal plane kinematics. J. Comp. Physiol. A Neuroethol. Sens. Neural Behav. Physiol. 195, 1011–1020.

Wild, A. L. (2004). Taxonomy and Distribution of the Argentine Ant, Linepithema humile (Hymenoptera: Formicidae). Annals of the Entomological Society of America 97, 1204–1215.

Wilkinson, E. J. and Sherk, H. A. (2005). The use of visual information for planning accurate steps in a cluttered environment. Behav. Brain Res. 164, 270–274.

Yarger, A. M. and Fox, J. L. (2016). Dipteran Halteres: Perspectives on Function and Integration for a Unique Sensory Organ. Integr. Comp. Biol. 56, 865–876.

Ye, S. and Comer, C. M. (1996). Correspondence of escape-turning behavior with activity of descending mechanosensory interneurons in the cockroach, Periplaneta americana. J. Neurosci. 16, 5844–5853.

Ye, S., Leung, V., Khan, A., Baba, Y. and Comer, C. M. (2003). The antennal system and cockroach evasive behavior. I. Roles for visual and mechanosensory cues in the response. J. Comp. Physiol. A Neuroethol. Sens. Neural Behav. Physiol. 189, 89–96.

Zill, S. N., Keller, B. R., Chaudhry, S., Duke, E. R., Neff, D., Quinn, R. and Flannigan, C. (2010). Detecting substrate engagement: responses of tarsal campaniform sensilla in cockroaches. J. Comp. Physiol. A Neuroethol. Sens. Neural Behav. Physiol. 196, 407–420.

Zollikofer, C. (1994). STEPPING PATTERNS IN ANTS - INFLUENCE OF SPEED AND CURVATURE. J. Exp. Biol. 192, 95–106.

Zurek, D. B. and Gilbert, C. (2014). Static antennae act as locomotory guides that compensate for visual motion blur in a diurnal, keen-eyed predator. Proc. Biol. Sci. 281, 20133072.

